# Hypoxia-directed tumor targeting of CRISPR/Cas9 and HSV-TK suicide gene therapy using lipid nanoparticles

**DOI:** 10.1101/2021.09.30.462477

**Authors:** Alicia Davis, Kevin V. Morris, Galina Shevchenko

**Affiliations:** Center for Gene Therapy, Beckman Research Institute, City of Hope, Duarte, CA, USA; Irell & Manella Graduate School of Biological Sciences, City of Hope, Duarte, CA, USA; Menzies Health Institute Queensland, School of Medical Science Griffith University, Gold Coast Campus, QLD 4222, Australia

**Keywords:** CRISPR/Cas9, HSV-tk, lipid nanoparticle, cancer, hypoxia

## Abstract

Hypoxia is a characteristic feature of solid tumors that contributes to tumor aggressiveness and is associated with resistance to cancer therapy. The hypoxia inducible factor-1 (HIF-1) transcription factor complex mediates hypoxia-specific gene expression by binding to hypoxia responsive element (HRE) sequences within the promoter of target genes. HRE driven expression of therapeutic cargo has been widely explored as a strategy to achieve cancer-specific gene expression. By utilizing this system, we achieve hypoxia-specific expression of two therapeutically relevant cargo elements: the Herpes Simplex Virus thymidine kinase (HSV-tk) suicide gene and the CRISPR/Cas9 nuclease. Using an expression vector containing five copies of the HRE derived from the vascular endothelial growth factor gene, we are able to show high transgene expression in cells in a hypoxic environment, similar to levels achieved using the CMV and CBh promoters. Furthermore, we are able to deliver our therapeutic cargo to tumor cells with high efficiency using plasmid packaged lipid nanoparticles (LNPs) to achieve specific killing of tumor cells in hypoxic conditions, while maintaining tight regulation with no significant changes to cell viability in normoxia.

## Introduction

Over recent years, many advances have been made in cancer therapy, including the development of targeted inhibitors and immunotherapies with greatly reduced adverse effects. Despite this, the current 5-year survival rate among all cancers is only 68%, dropping to under 10% for certain cancer types, including pancreatic cancer and glioblastoma (1). The resistance of these high-risk cancers to current therapies begets a need to develop novel treatment modalities to improve patient survival and quality of life. Several unique cancer treatment strategies have emerged by repurposing proteins found in viruses and bacteria to promote cell death, including the Human Simplex Virus thymidine kinase (HSV-tk) suicide gene and the CRISPR/Cas9 nuclease. Although both of these proteins have shown promise as a treatment option for various cancers, there is a growing need to direct tumor-specific expression of these proteins in order to minimize the risk of off-target toxicity and, in the case of Cas9, off-target mutations.

The hypoxic microenvironment offers a unique opportunity to target these exogenous proteins to regions of the tumor where the most aggressive and treatment resistant cancer cells often reside. Solid tumors account for approximately 90% of all human malignancies (2). As a result of vascular abnormalities that lead to low intratumoral blood flow, up to 50-60% of locally advanced solid tumors develop areas of low O_2_ (<10 mmHg) partial pressure compared to their surrounding tissues (3–5). This hypoxic state has been associated with increased tumor aggressiveness and resistance to current therapies (6). Of all the proteins induced by hypoxic conditions, hypoxia-inducible factors (HIF) and their downstream targets are the most well studied. Under normoxic conditions, the protein expression of HIF1α is short-lived, with a half-life of approximately 5 minutes, as a result of rapid ubiquitin-dependent proteasomal degradation (7,8). Once induced, HIF heterodimers bind to hypoxia response elements (HRE) within the promoters of target genes, including vascular endothelial growth factor (VEGF). When gene expression is driven by promoters containing multiple copies of HRE sequences, hypoxia-specific gene expression is obtained. This HRE-directed gene expression has been well established in multiple systems, including hypoxia-specific CAR-T cells and multiple other therapies (9,10). While such hypoxia-specific gene expression has been explored for the HSV-tk suicide gene system (11)(12), we aimed to improve on the existing systems and delivery methods. In addition to this, hypoxia mediated regulation has never been applied to the CRISPR-Cas9 system to achieve specific expression of the Cas9 protein.

One of the greatest challenges for nucleic acid based therapeutics is the method of delivery. Delivery of HSV-tk and CRISPR-Cas9 *in vivo* has been demonstrated with varying levels of success using adeno-associated viral (AAV) vectors (13–15). However, significant limitations exist for AAV mediated delivery of CRISPR-Cas9 stemming from the large size of the Cas9 system and promoters necessary to direct replication in specific cell types (16). Additionally, the application of AAV therapies remains limited based on both innate and adaptive immune responses (17). Lipid nanoparticles (LNPs) are an emerging alternative to AAVs (18) that offer a protective, flexible, and simple platform to encapsulate and protect therapeutic cargo, including drugs, siRNAs, mRNA, protein and DNA (19–23). While many efforts are being made in order to increase the specificity of LNP mediated delivery to tumor cells, LNPs are known to commonly deliver cargo to the liver, spleen and kidneys, which becomes a major cause of concern for off-target activity (24,25). Based on these concerns, we proposed to direct conditional expression of HSV-tk and Cas9 to increase LNP specificity and limit potential off-target toxicity.

In the present study, we report on the development of both HSV-tk suicide gene and CRISPR-Cas9 DNA lipid nanoparticle based therapeutic systems that enable hypoxic tumor specific expression, in order to minimize concerns regarding off-target toxicities. To this end, we expressed these protein systems from a minimal promoter in combination with five copies of the hypoxia responsive element derived from the human VEGF enhancer. Through extensive testing using both hypoxia drug mimetics and true hypoxia, we have observed tightly regulated hypoxia-dependent cancer cell death using two distinct gene therapy approaches.

## Materials and Methods

### Cell Lines and Cell Culture

U251 cells were obtained as a kind gift from the lab of Dr. Stephen Forman (City of Hope, CA). The human cell lines 293FT (Invitrogen™ R70007), HT-1080 (ATCC® CCL-121™), NCI-H1299 (ATCC® CRL-5803™), and HeLa (ATCC® CCL-2™) cells were cultured according to the respective guidelines in a humidified incubator with 5% CO2 at 37°C. The following media and supplements were used in this study for routine tissue culture: DMEM (Corning® 10-017-CV), RPMI (Corning® 10-040-CV), EMEM (Corning® 10-009-CV), Benchmark™ FBS (GeminiBio 100-106), DPBS (Corning® 21-040-CMR), 0.25% Trypsin-EDTA (Gibco™ 25200056). All cell lines were routinely tested for mycoplasma contamination using the MycoAlert™ PLUS Mycoplasma Detection Kit (Lonza LT07-710).

### Treatment of Cells

#### Transfection

Plasmid transfection was carried out using Lipofectamine™ 3000 (Invitrogen™ 11668019) or ViaFect™ (Promega™ E4981) according to the protocol provided by the manufacturer. A complete list of plasmids used in this study can be found in **Supplemental Table 1**.

#### Hypoxia

For chemical induction of hypoxia, cells were treated with 10μM of the HIF prolyl-hydroxylase inhibitor BAY 85-3934(26) (Molidustat) (Cayman Chemical 15297). The BAY 85-3934 inhibitor was dissolved in DMSO (ATCC® 4-X™) at a stock concentration of 10mM. For hypoxia treatment, cells were placed in a BD GasPak™ EZ anaerobe pouch (BD 260683) for the times indicated in the figure legends.

#### Ganciclovir treatment

For experiments involving the suicide gene HSV-tk, cells were treated with ganciclovir (ACROS Organics™ 461710010) at the indicated concentrations. The ganciclovir stock solution was made by dissolving the drug in DMSO at a final concentration of 100mM.

### Cell Viability Assay

Cell viability was assessed using a 10X Alamar reagent: 650μM rezasurin (Sigma Aldrich® 199303), 78μM methylene blue (Sigma Aldrich® M9140), 1mM potassium ferrocyanide (ACROS Organics™ 196781000), and 1mM potassium ferricyanide (ACROS Organics™ 424130050) in DPBS, diluted to a final concentration of 1X in the cell culture medium (U.S. Pat. No. 5501959). After addition of the Alamar reagent, cells were incubated in a humidified incubator with 5% CO2 at 37°C for 1-6 hours before reading the fluorescence of the solution using the Promega GloMax® Discover Microplate Reader (Ex 525 nm: Em 580-640 nm).

### Luciferase Activity Assay

Firefly Luciferase activity was detected using the Luciferase Assay System (Promega E4550) according to the protocol supplied by the manufacturer. Luminescence was detected using the Promega GloMax® Discover Microplate Reader.

### T7 Endonuclease Assay

To survey genomic targeting activity, genomic DNA was extracted from cells at the indicated time points using the Maxwell® RSC Cultured Cells DNA Kit (Promega AS1620). DNA was amplified using Q5® High-Fidelity 2X Master Mix (New England BioLabs® M0492L) from 100ng of genomic DNA according to the protocol provided by the manufacturer. The following primers (sequences 5’-3’) were used for genomic amplification of the PLK1 locus: PLK1-Fwd GGTTTGGTTTCCCAGGCTATC, PLK1-Rev TCCAGAAGCTGACTTCCCTATC. After PCR amplification, 100-200ng of PCR product was denatured and re-annealed in a thermal cycler: 3 minutes at 98°C then cooled to 25°C using a ramp rate of 0.1°C/second. After re-annealing the DNA, 5U of T7 Endonuclease I (New England BioLabs® M0302L) was added and the reaction was incubated at 37°C for 20 minutes. After completion, the reaction was loaded onto an agarose gel for visualization of cleavage efficiency. Cleavage efficiency was determined using the following calculation: percent cleavage = 100 × (1−(1−fraction cleaved)^½^), where fraction cleaved = (cleavage products)/(cleavage products + parental band).

### Western Blotting

For western blotting, cells were lysed in 1X RIPA lysis buffer: 50mM Tris HCl (pH 7.4), 150mM NaCl, 1.0% (v/v) NP-40, 0.5% (v/v) sodium deoxycholate, and 0.1% (v/v) SDS. To facilitate lysis, the lysate was then sonicated on the high setting for a total of three 30 second cycles using the Bioruptor® Plus (Diagenode Inc.). For SDS-PAGE electrophoresis, 20-30 μg of protein was loaded per well. The proteins were then transferred to a nitrocellulose membrane using the Trans-Blot Turbo Transfer System (Bio-Rad Laboratories). To enhance detection of HIF1α, the membrane was treated using the Pierce™ Western Blot Signal Enhancer (Thermo Fisher Scientific 21050) before blocking. Antibodies were diluted in blocking solution: 2% bovine serum albumin in 1X Tris-Buffered Saline with 0.1% (v/v) Tween-20. Protein detection was performed using the SuperSignal™ West Pico PLUS Chemiluminescent Substrate (Thermo Fisher Scientific) using the ChemiDock system (Bio-Rad Laboratories). The following antibodies were used in this study: anti-GAPDH (1:10,000; Santa Cruz Biotechnology sc-47724), anti-HIF1α (1:1000; Bethyl Laboratories A700-001), anti-spCas9 (1:10,000; Sigma-Aldrich SAB4200701), anti-thymidine kinase (1:2000; Invitrogen™ PA5-67984), anti-Histone H3 (1:10,0000; Cell Signaling Technologies 14269), anti-mouse IgG HRP conjugate (1:10,000; Bio-Rad Laboratories 1706516), anti-rabbit IgG HRP conjugate (1:10,000; Bio-Rad Laboratories 1706515).

### Brightfield and Fluorescence Microscopy

Induction of HRE-eGFP for each experiment was verified by visualizing cells using brightfield and fluorescence microscopy. Images were captured using the ZEISS Axio Vert A.1 inverted microscope. Images were processed for export using ZEN Blue v2.6 software (ZEISS).

### Immunofluorescence

#### γH2AX DNA damage staining

Cells were grown using the Nunc™ Lab-Tek™ II Chamber Slide™ System (Thermo Scientific™ 154534PK) pre-treated with a 0.01% Poly-L-Lysine solution (Sigma Aldrich® A-005-M) to facilitate cell adhesion. Cells were transfected the next day and treated with either DMSO vehicle or 10μM BAY 85-3934. After 48 hours, cells were fixed for 10 minutes at room temperature using a 4% paraformaldehyde solution (Thermo Fisher Scientific AAJ19943K2). The fixed cells were then permeabilized using a 0.25% Triton-X solution and blocked using 1% BSA w/v in PBS for 1 hour at room temperature. Primary antibody incubation was performed overnight at 4°C, while secondary antibody incubation was performed for 1 hour at room temperature. The following antibodies were used: γH2AX (1:1000; Invitrogen™ MA12022), HSV-tk (1:400; Invitrogen™ PA5-67984), Alexa Fluor 488 conjugated Goat anti-Mouse IgG (1:1000; Invitrogen™ A28175), Alexa Fluor 555 conjugated Goat anti-Mouse IgG (1:1000; Invitrogen™ A21422), and Alexa Fluor 647 conjugated Goat anti-Rabbit IgG (1:1000; Invitrogen™ A27040). After staining, the slides were mounted using ProLong™ Glass Antifade Mountant (Invitrogen™ P36982) and sealed using generic brand nail polish. Staining was visualized using the ZEISS LSM 880 with Airyscan confocal microscope. Images were processed for export using ZEN Blue v2.6 software (ZEISS).

### Flow Cytometry

Following transduction or transfection, cells were trypsinized, washed and resuspended in complete growth medium and placed on ice. Flow cytometry was conducted using the BD Accuri™ C6 Cytometer (BD Biosciences). Data was analyzed using FlowJo™ v10.8 software (BD Biosciences).

### Lipid Nanoparticle Assembly and Analysis

#### Lipids

The following lipids 1-stearoyl-2-oleoyl-*sn*-glycero-3-phosphocholine (SOPC), cholesterol, and 1,2-dimyristoyl-rac-glycero-3-methoxypolyethylene glycol-2000 (PEG-DMG) were purchased from Avanti Polar Lipids (Alabaster, AL, USA). 2,2-dilinoleyl-4-(2-dimethylaminoethyl)-[1,3]-dioxolane (DLin-KC2-DMA) was purchased from (MedChemExpress; Monmouth Junction, NJ, USA). Lipophilic dye DiL (DilC_18_(3) (1,1’-dioctadecyl-3,3,3’,3’-tetramethylindocarbocyanine perchlorate) at 1mM stock in ethanol (Invitrogen; Carlsbad, CA, USA) was used to label nanoparticles to observe *in vitro* delivery. Lipids were dissolved in absolute ethanol, aliquoted in amber glass vials, and stored at -20°C.

#### LNP Synthesis

LNPs were prepared as described in Kulkarni *et*.*al. (23)*. Lipids were prepared at a 50:10:39:1 (KC2:SOPC:Chol:PEG) molar ratio. Lipids in ethanol were mixed with plasmid prepared in an aqueous phase of 100mM sodium citrate pH 4 at a mol cationic lipid: mol DNA (N:P) ratio of 6:1 using the NanoAssemblr Benchtop machine (Precision NanoSystems; Vancouver, BC, Canada). This machine contains a microfluidic chip by which the injected lipids and nucleic acids are rapidly mixed in a staggered herringbone pattern at a total flow rate of 12mL/min. The controlled mixing of the aqueous and organic streams produces homogeneous nanoparticles. Immediately following the mixing process, the nanoparticles were diluted 1:4 in 1X PBS to reduce the amount of ethanol present in solution. The nanoparticle solution was further diluted with 1X PBS up to 15 mL and then concentrated using a 10 kDa Amicon ultra-15 filter (Millipore; Burlington, MA, USA) via centrifugation at 2,000 x G for 30 minutes. The column flow through was discarded and another 15 mL 1X PBS was added to the nanoparticles which were centrifuged at 2,000 x G for 30 mins. This step was repeated one additional time for a total of 3 washes. The concentrated nanoparticles were then pushed through a 0.22μm filter and stored at 4°.

#### LNP Characterization

To measure the amount of plasmid encapsulated inside the nanoparticles the Quant-IT Picogreen assay was carried out (P7589, Molecular Probes; Eugene, OR, USA). The standard protocol was modified to include a 15-minute, 37°C incubation of the nanoparticles in the presence of 2% Triton to facilitate release of the encapsulated nucleic acids. %encapsulation = (DNA-LNP in 2%Triton - DNA-LNP in TE)/DNA-LNP in 2%Triton-X100 (based on methods in (27).

Size and polydispersity of the nanoparticles was evaluated using dynamic light scattering (DLS) analysis and relative surface charge was determined by measuring zeta potential on a ZetaPals instrument (Brookhaven Instruments Corporation; Holtsville, NY, USA). Samples were diluted in distilled water for both DLS and zeta potential measurements. Size and polydispersity were evaluated in five runs of 1 minute and 30 second duration, while zeta potential was evaluated in 10 runs of 30 cycles/run. In order to visualize the nanoparticles, negative staining electron microscopy was carried out. Nanoparticles were diluted 1/10 in 1X PBS and absorbed to glow-discharged, carbon-coated 200 mesh EM grids. Conventional negative staining was carried out using 1% (w/v) uranyl acetate and images were captured using an FEI Tecnai 12 transmission electron microscope and Gatan OneView CMOS camera.

### Lentiviral generation of stable cells

Lentiviral production was performed in 293FT cells as described in Urak *et*.*al*. (28). HeLa cells were transduced at MOI = 0.2 using lentiviral particles containing a pLenti-EF1a-eGFP-ffluc-BSR transfer plasmid. Selection for stably transduced cells was performed by adding blasticidin (Gibco™ A1113903) to a final concentration of 5 μg/mL.

### Statistical Analysis

All data are presented as mean ± S.D. and are representative of a minimum of two independent experiments. Statistical analysis was performed using Prism v.9.1.2 (GraphPad). Comparisons of three or more groups were analyzed using one-way or two-way ANOVA tests, when appropriate. One-way ANOVA tests were corrected for multiple comparisons using the Dunnett correction method, while two-way ANOVA tests were corrected using the Tukey method. P-value < 0.05 was considered statistically significant.

## Results

### Development and characterization of a hypoxia regulated protein expression system

In order to determine the best combination of regulatory elements for hypoxia-specific gene expression, we conducted a luciferase screen in HeLa cells in the presence or absence of the hypoxia mimetic BAY 85-3934 (Figure S1A). BAY 85-3934 is a potent small-molecule inhibitor of HIF prolyl hydroxylases that stabilizes HIF1α from proteasomal degradation (26). We confirmed its ability to stabilize HIF1α using a HRE eGFP reporter (Figure 1A) and were able to see induction of HIF1α protein by western blot to levels similar to those achieved in true hypoxia (Figure 1B). We designed fusion constructs consisting of firefly luciferase combined with regions of the HIF1α oxygen-dependent degradation domain (ODD) that were designated as either short (residues 344-417 of HIF1α) or long (residues 402-603 of HIF1α). These regions were chosen as they encompass the two key proline residues (Pro402 and Pro564) of HIF1α that are important for proteasomal degradation of the protein under normoxic conditions (29). In order to determine the contribution of promoter context to the expression of our fusion constructs, we expressed these constructs from three different promoters: (A) cytomegalovirus (CMV) promoter, (B) human phosphoglycerate kinase (PGK) promoter, and (C) a minimal CMV promoter combined with five copies of the human VEGF HRE (5HRE) (30). As the 5HRE promoter provided superior control of gene expression under hypoxic conditions for all of the constructs (Figure S1A), we proceeded to examine its contribution to the expression of the luciferase fusion constructs in HeLa cells (Figures 1C and 1D) and H1299 cells (Figures S1B and S1C). While the longODD construct offered the best induction of protein expression in hypoxia when driven from the HRE promoter (900-fold induction in HeLa cells), it limited overall protein expression when compared to levels that were obtained from a CMV promoter. Consequently, protein expression from the 5HRE promoter in the absence of any ODD fusion provided the best combined regulation and expression under hypoxic conditions.

**Figure 1.**
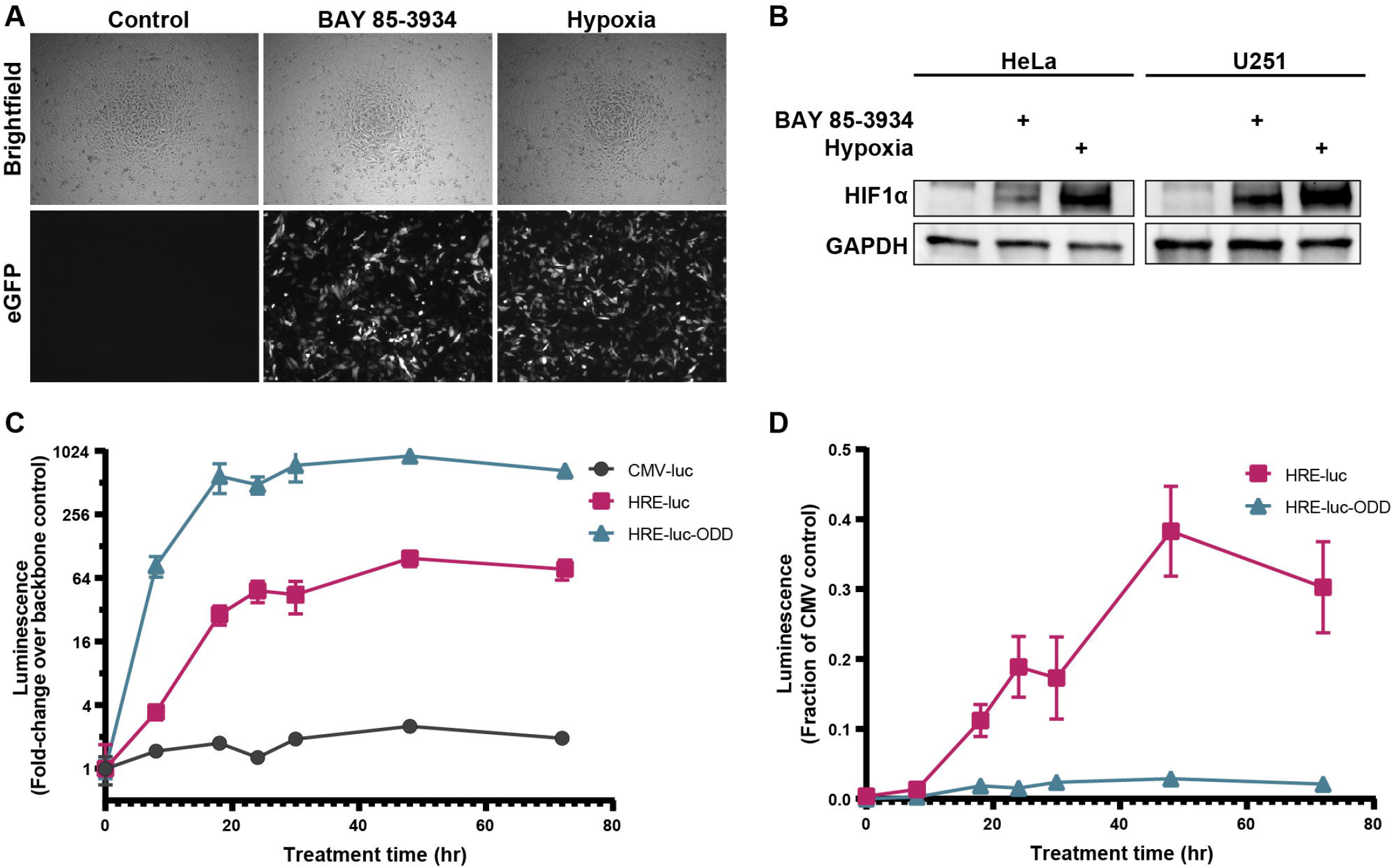
Optimization of Hypoxia-Regulated Luciferase Reporter Systems. (A) Representative images of 5HRE eGFP-expressing HeLa cells grown under control conditions, in the presence of 10μM BAY 85-3934, or in hypoxia. (B) Western blot showing expression of HIF1α in HeLa and U251 cells grown for 6 hours under control conditions, in the presence of 10μM BAY 85-3934, or in hypoxia. (C) Time-course induction of normalized luciferase activity relative to the respective backbone control in HeLa cells expressing the indicated constructs grown in the presence of 10μM BAY 85-3934, or vehicle control (0.1% v/v DMSO). (D) Same as in (C), except the luciferase signal is normalized to the CMV backbone. Data are represented as mean ± S.D of triplicate treated samples.

### Hypoxia-specific expression of HSV-tk

We initially tested the HRE-driven system with the well-established suicide gene HSV-tk. This suicide gene has been widely explored as an anti-cancer therapeutic in combination with acyclic guanosine analogs such as ganciclovir (GCV). GCV is largely inert in the absence of HSV-tk as it cannot be used as a substrate by cellular enzymes (31). In the presence of HSV-tk however, GCV becomes phosphorylated by the viral protein to produce GCV-monophosphate, a form of the drug that can now be further processed by cellular enzymes to form GCV-triphosphate (32). Incorporation of GCV-triphosphate into nascent DNA strands subsequently causes chain termination during DNA synthesis, leading to cell death (33). Notably, cells expressing HSV-tk can transfer GCV-triphosphate to neighboring cells through gap junctions and apoptotic vesicles, which can then induce cell death in HSV-tk negative cells through a phenomenon known as the bystander effect (32).

This suicide gene system has been optimized through the testing of HSV-tk mutants and fusion proteins (34,35). Previous studies identified a HSV-tk variant (mutant 30) derived by random mutagenesis that displayed enhanced sensitivity to GCV and enhanced the bystander effect when compared with wild-type HSV-tk *in vivo* (36). Based on these findings, we expressed wild-type HSV-tk and mutant 30 from the 5HRE promoter (Figure 2A). We confirm HRE-specific induction of both HSV-tk constructs at the protein level in both BAY 85-3934 and hypoxia treated Hela and U251 cells (Figure 2B and Figure S2A). In the absence of the hypoxia mimetic or hypoxia treatment there is no detectable protein expression of HSV-tk from the HRE constructs. Interestingly, HRE-HSV-tk WT protein expression under hypoxia is at nearly the same level as the CMV counterpart. Cells transfected with CMV HSV-tk WT or mutant 30 display reduced viability as expected when treated with GCV (Figure 2C and Figure S2B-C). However, cells expressing HRE directed HSV-tk WT or mutant 30 only exhibit reduced cell viability when cultured in the presence of GCV and the hypoxia mimetic BAY 85-3934 or true hypoxia (Figure 2C). In order to visualize the hypoxic induction of HSV-tk mediated DNA damage occurring in these cells we performed γH2AX staining. Indeed, we observe γH2AX staining in cells receiving HRE-HSV-tk mutant 30 during BAY 85-3934 treatment but do not observe γH2AX staining in the respective DMSO control (Figure 2D). Collectively, these data demonstrate that HRE driven expression of HSV-tk WT or mutant 30 is tightly regulated and significantly reduces cell viability only during hypoxic conditions.

**Figure 2.**
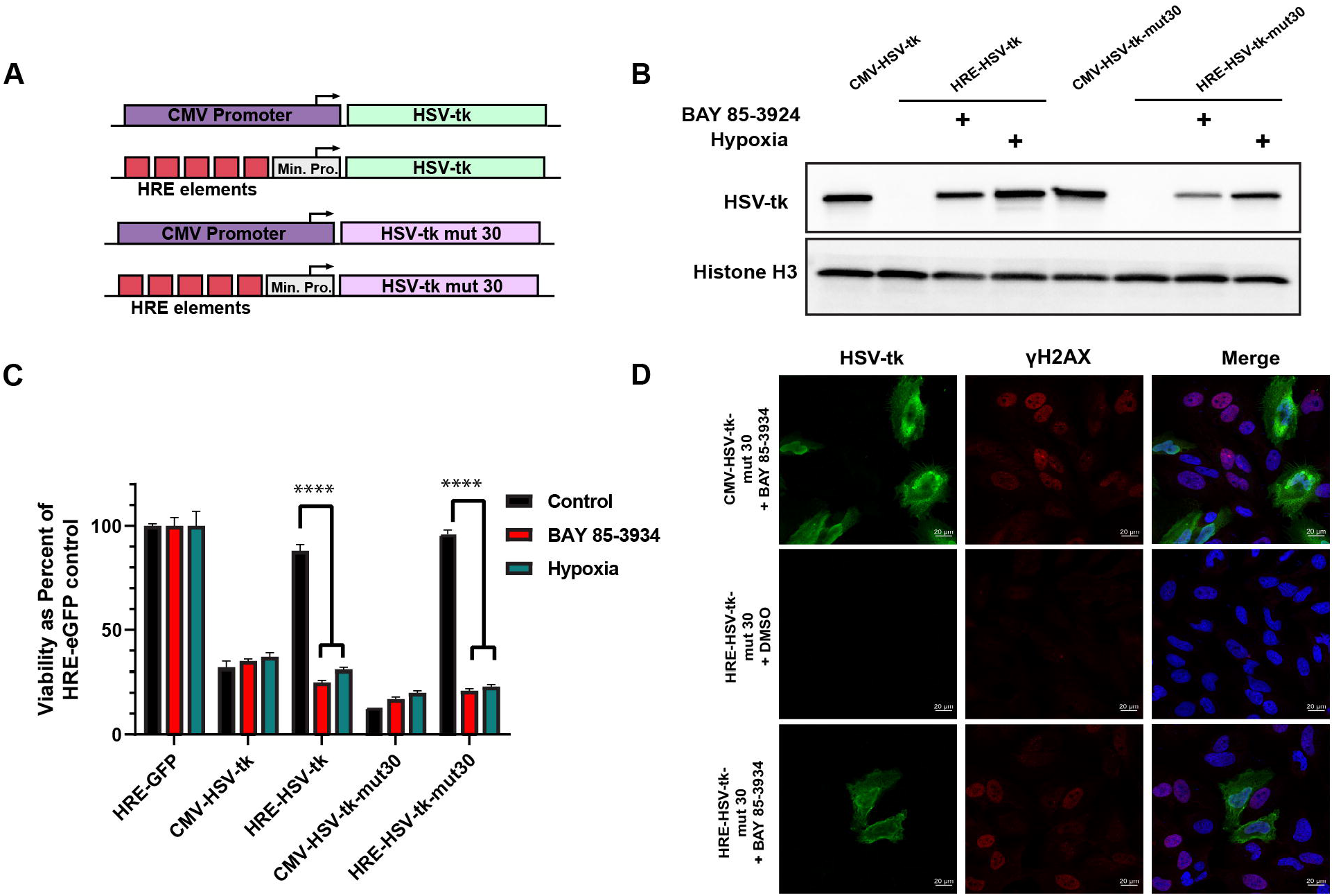
Hypoxia-regulated expression and activity of HSV-tk. (A) Schematic depicting the two promoters used to direct expression of HSV-tk constructs. The cytomegalovirus (CMV) promoter drives the constitutive expression of HSV-tk thereby serving as the positive control for these experiments. The hypoxia-regulated promoter consists of a minimal CMV promoter and five copies of the human VEGF HRE. (B) Western blot analysis of HeLa cells transfected with either the CMV or HRE-driven HSV-tk WT or mutant 30 constructs. Transfected cells were cultured in the presence of 10 μM BAY 85-3934 or hypoxia for 24 hours prior to harvesting for protein analysis. (C) An Alamar assay for viability was carried out at 120 hours post-transfection of the indicated HSV-tk constructs in HeLa cells cultured in 1μM GCV with either 10μM BAY 85-3934 or hypoxia. Cells undergoing hypoxia treatment were exposed to two intermittent periods of hypoxia for 24 hours each. (D). Confocal fluorescent microscopy for DNA damage signaling marker γH2AX during HSV-tk treatment. HeLa cells were transfected with the HRE-HSV-tk mutant 30 construct and cultured in the presence of 1μM GCV with either 10 μM BAY 85-3934 or DMSO vehicle control for 48 hours. Cells were stained for HSV-tk (green), DNA damage marker γH2AX (red), and the nuclear morphology indicator Hoescht 33342 (blue). Scale bar=20μm. Data are represented as mean ± S.D of triplicate treated samples; ****p<0.0001.

### Hypoxia-specific expression of CRISPR/Cas9

The Cas9 protein is a repurposed prokaryotic RNA-guided dsDNA nuclease used for mammalian gene editing that can be programmed to cut any genomic locus. CRISPR-Cas9 offers a unique opportunity to treat cancer by permanently disrupting genes essential for tumor cell survival. However, there is currently no reliable way to specifically deliver and express the CRISPR-Cas9 nuclease in cancer cells. To overcome this limitation, many efforts have focused on targeting unique sequence variants that are found only in cancer cells, such as viral and fusion oncogenes (37–40), minimizing the risk of any potential toxicity in normal tissue. Nevertheless, in addition to the largely limited number of targets that can be pursued with this approach, there is also a growing concern about the risks of off-target mutations that can arise with unchecked expression of the Cas9 throughout the body (41–43). Based on these concerns, we set out to evaluate the potential of an HRE-directed CRISPR/Cas9 to specifically induce cancer cell death in hypoxia.

To determine the best gRNA targets, we screened gRNAs targeting polo-like kinase 1 (PLK1) and the consensus sequence of AluY elements for their ability to suppress cancer cell growth. Polo-like kinase 1 (PLK1) has been extensively evaluated as both an siRNA and Cas9 target to induce apoptosis in a broad range of cancer cells (44,45). Consistent with previous work, all gRNAs targeting PLK1 suppressed cell viability, with gRNA-3 being used in all subsequent experiments as it appeared to be the most potent in multiple cell lines (Figures 3A and S3A). A gRNA against the consensus AluY sequence was further included as it has the potential to induce thousands of dsDNA breaks throughout the genome, leading to a significant reduction in cell viability (Figures 3A and S3A).

**Figure 3.**
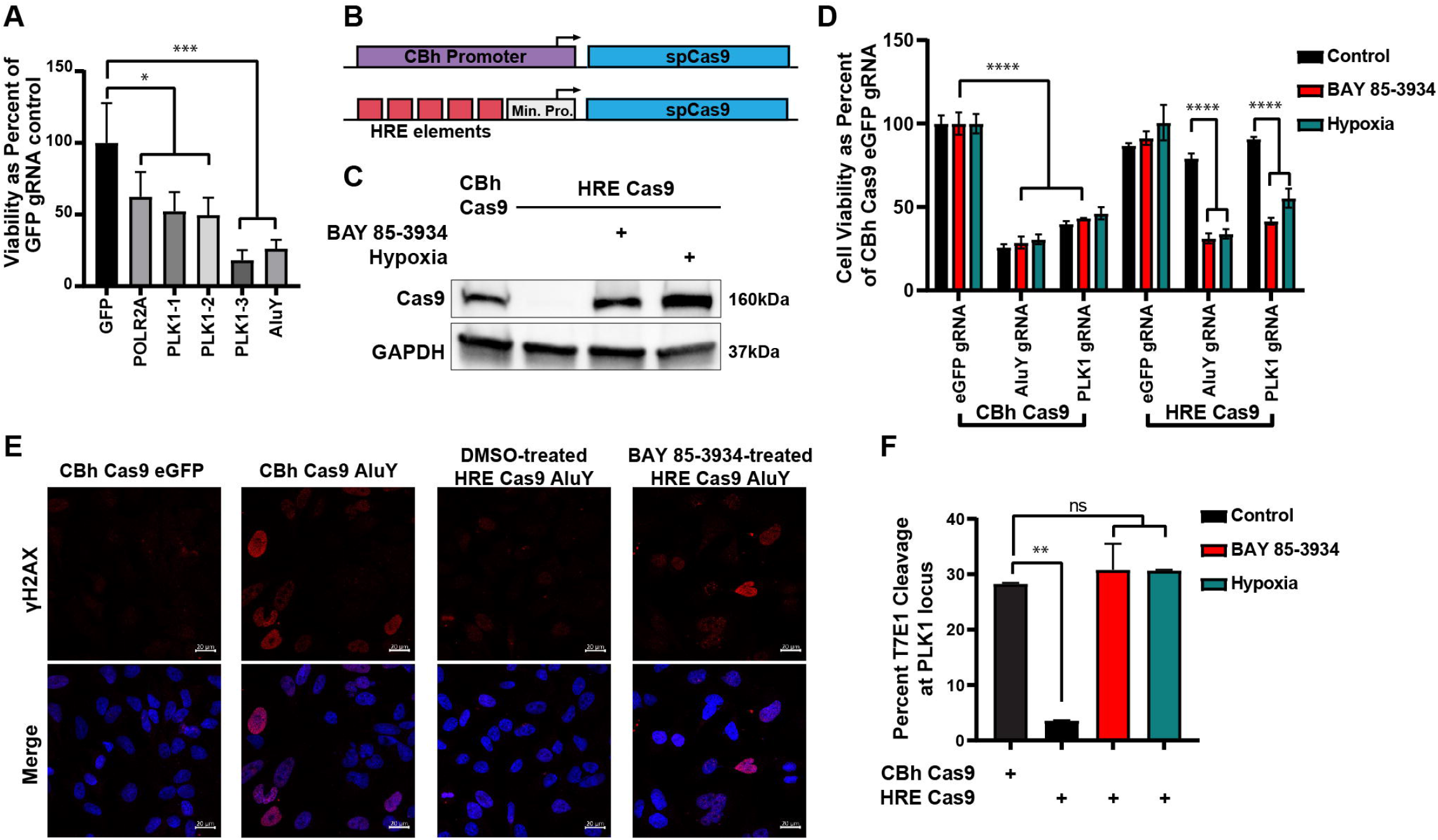
Hypoxia-regulated expression and activity of CRISPR/Cas9. (A) Alamar assay looking at the viability of HeLa cells 72 hours post-transfection with a CBh-driven Cas9 and the indicated gRNAs. (B) Schematic of the two spCas9 DNA constructs used in this study. The ubiquitously expressed Cas9 nuclease is driven from the CBh promoter (CMV early enhancer, chicken β-actin promoter, and a chimeric chicken β-actin/MVM intron), while the hypoxia-regulated Cas9 is driven from a minimal promoter combined with five copies of the human VEGF HRE. (C) Western blot showing expression of Cas9 in HeLa cells transfected with either the CBh- or HRE-driven Cas9. HeLa cells transfected with the HRE-driven Cas9 were cultured in the presence of 10μM BAY 85-3934 or hypoxia for 24 hours before harvesting lysates. (D) Alamar assay comparing viability of HeLa cells 120 hours post-transfection with the indicated constructs and treated with the indicated conditions. Cells were cultured in the presence of the hypoxia mimetic, BAY 85-3934, for the entire course of the experiment. For hypoxia-treatment, cells were treated with intermittent hypoxia. Cells were placed into hypoxic conditions for a total of 48 hours, in 24 hour intervals. (E) Confocal fluorescent microscopy of HeLa cells 48 hours post-transfection with the indicated constructs. Cells were stained with γH2AX (red) to indicate DNA damage and Hoechst 33342 (blue) to indicate nuclear morphology. Scale bar = 20μm. (F) T7 endonuclease I assay measuring Cas9-mediated cleavage at the PLK1 locus in HeLa cells 48 hours post-transfection with the indicated constructs and treated in the indicated conditions. Data are represented as mean ± S.D of triplicate treated samples, except (F) which represents mean ± S.D of two independent samples; n.s. not significant; *p<0.05; **p<0.01; ***p<0.001; ****p<0.0001.

To determine the feasibility and efficacy of a hypoxia-regulated Cas9 expression system, we transfected cells with the Cas9 nuclease expressed from either the CBh or the 5HRE promoter (Figure 3B). We observed HRE-specific expression of the Cas9 nuclease in cells cultured with the hypoxia mimetic BAY 85-3934 or in hypoxia, with almost undetectable levels of expression in control conditions (Figure 3C and S3B). We saw a significant reduction in cell viability in cells transfected with the AluY and PLK1 gRNAs when the Cas9 nuclease was expressed from the CBh promoter in all conditions (Figure 3D and S3C). However, a significant reduction in cell viability was only observed when cells were treated in hypoxic conditions when these gRNAs were expressed in the presence of the HRE-driven Cas9 (Figure 3D and S3C). For cells treated with the AluY gRNAs, we observed an increase in DNA damage using the HRE driven Cas9 nuclease only in hypoxic conditions, while there was evidence of DNA damage in both hypoxic and normoxic conditions using the CBh driven Cas9 (Figure 3E). Next, we wanted to confirm that Cas9 cleavage was only observed in hypoxic conditions when the expression of the nuclease is driven from the HRE promoter. To this end, we utilized a reporter cell line containing an eGFP firefly luciferase fusion protein. Cas9 cleavage with our eGFP gRNA results in the disruption of the fusion protein in these cells, giving us an opportunity to use the expression of eGFP and luciferase as an indicator of Cas9 activity. We observed a specific reduction in luciferase signal in a hypoxia regulated fashion for cells transfected with the HRE regulated Cas9 construct (Figure S3D). However, as this was not a direct measure of Cas9 cleavage activity, we performed a T7 Endonuclease I cleavage assay in order to directly survey genomic editing at the PLK1 locus. We were able to show hypoxia regulated Cas9 mediated editing at the PLK1 locus, with minimal cleavage at this locus in normoxic conditions (Figure 3F and S3E). Overall, we demonstrate a HRE promoter driven Cas9 system that is capable of specifically reducing the viability of hypoxic cancer cells.

### Delivery using LNPs

As our method for achieving hypoxia-specific expression for HSV-tk and Cas9 requires transcriptional activation at the promoter level, we needed a vehicle that could facilitate delivery of a DNA-based system in order to achieve regulated expression of these genes. While AAV has been widely used as a vector for gene therapy to deliver DNA sequences to target cells, the large size of the spCas9 nuclease together with its gRNA (∼5kb) makes it largely incompatible with an AAV-based delivery vehicle due to the ∼4.7kb packaging capacity of AAVs. In light of this, we decided to deliver these transgenes using plasmid packaged lipid nanoparticles (LNPs). Recently, the feasibility of LNP therapies have gained immense traction with the success of the mRNA based COVID-19 vaccines produced by Moderna and BioNTech/Pfizer (46). LNPs are typically formulated using a mixture of four lipids: (1) an ionizable cationic lipid to facilitate cargo encapsulation and intracellular delivery, (2) a PEG containing lipid to modulate particle size, reduce aggregation and increase circulation time, (3) cholesterol to enhance particle stability, particularly in circulation, and (4) neutral helper lipids to further promote formulation stability of the lipid bilayer (47,48). In order to deliver DNA, we utilized the formulation optimized by Kulkarni *et al*., 2018 in which the ionizable lipid DLin-KC2 demonstrated delivery of a Firefly luciferase or GFP plasmid. In order to test the feasibility of delivering a Cas9 plasmid, we encapsulated a 9.3kb Cas9-T2A-eGFP plasmid into LNPs using a nitrogen to phosphate (N:P) ratio of 6. Flow cytometry analysis of eGFP expression 48 hours after LNP treatment demonstrates efficient delivery of the plasmid into cells (Figure S4A). Furthermore, these LNPs are well tolerated as all cell lines tested remain greater than 90% viable 48 hours after treatment (Figure S4B).

Based on the successful delivery of the Cas9-T2A-eGFP plasmid using the KC2 DNA-LNP formulation, we moved forward and synthesized LNPs for each DNA construct used in this study. These LNPs are approximately 136±10.4 nm in size, with a low polydispersity of 0.09±0.03, and a zeta potential of 21.7±5.6 mV (Figures 4A and S4C). Visualization of these nanoparticles was carried out using transmission electron microscopy and revealed a consistent spherical morphology (Figure S4D). We first evaluated induction of GFP expression in HeLa cells following addition of LNPs packaged with the HRE-eGFP plasmid through both fluorescent microscopy and flow cytometry. We observe efficient uptake of Cy3-labeled HRE-eGFP LNPs in all cells but only observe GFP expression in cells cultured in BAY 85-3934 or hypoxia (Figure 4B). Flow cytometry analysis of HRE-eGFP LNP treated HeLa cells supports our microscopy observations and shows strong induction of GFP in cells cultured in BAY 85-3934 or hypoxia (Figure 4C).

**Figure 4.**
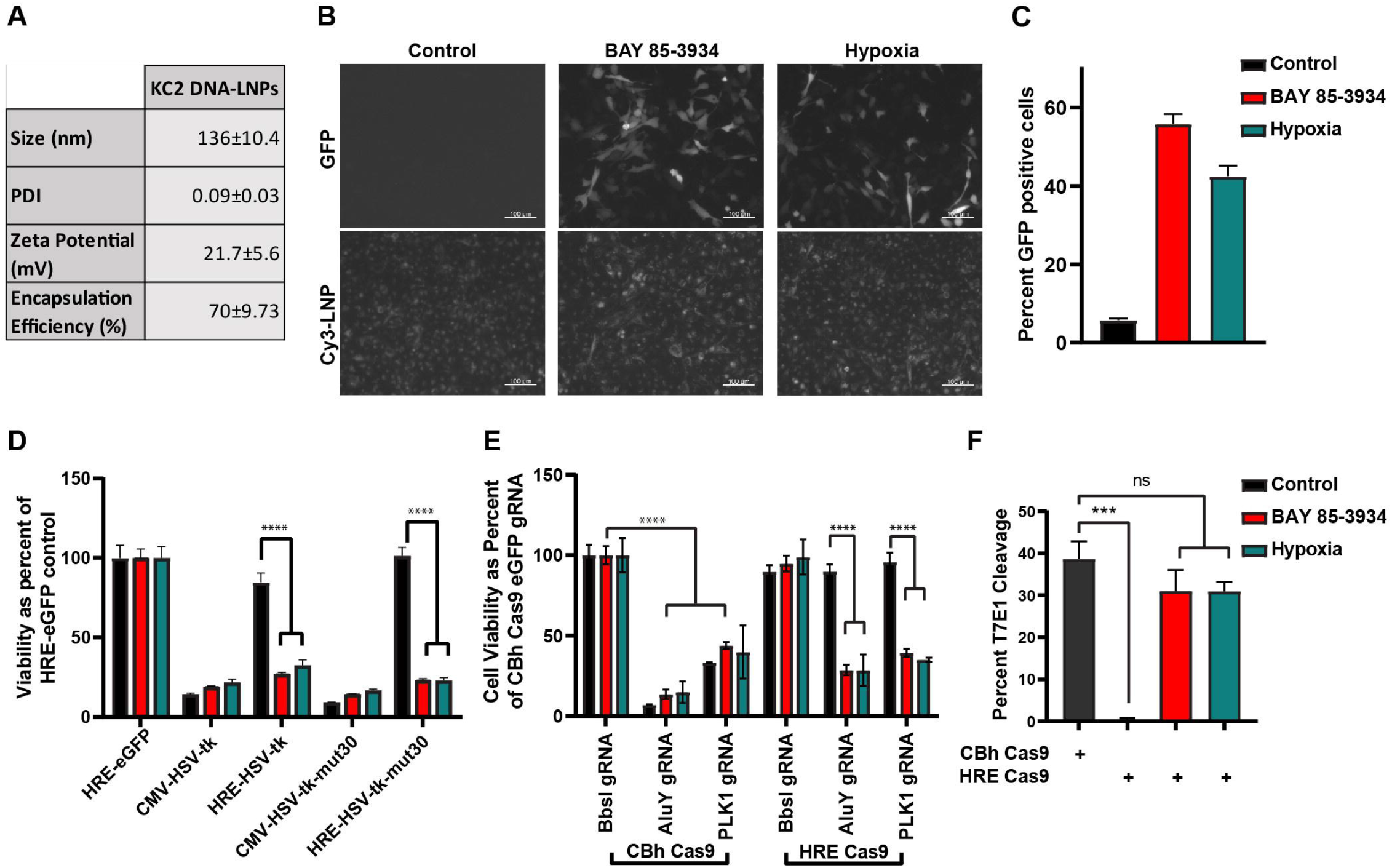
Delivery of Hypoxia-regulated HSV-tk and CRISPR/Cas9 using LNPs. (A) Lipid nanoparticles encapsulating our HSV-tk and Cas9 constructs were synthesized and characterized for size, polydispersity, zeta potential, and encapsulation efficiency. Data represented as mean ± S.D; for n=13 LNPs each containing different DNA cargo. (B) Fluorescent microscopy images of eGFP induction (top panel) 48 hours after LNP addition. HeLa cells received HRE-eGFP packaged LNPs and were treated with either DMSO control, 10μM BAY 85-3934, or cultured in hypoxia for 24 hours. LNPs used in this study were labeled with Cy3 allowing for visualization of uptake in cells (bottom panel). Scale bar=100μm. (C) Flow cytometry analysis of eGFP expression in HeLa cells 72 hours after addition of HRE-eGFP packaged LNPs. Cells were treated with either DMSO control, 10μM BAY 85-3934, or cultured in hypoxia for 24 hours. (D) Alamar assay evaluating potency of LNPs delivering the indicated HSV-tk constructs to HeLa cells in the presence of 1μM GCV and DMSO, BAY 85-3934, or hypoxia treatment at 120 hours post LNP treatment. (E) Alamar assay comparing viability of HeLa cells 120 hours post addition of LNPs packaged with the indicated constructs. Cells were cultured in the presence of the hypoxia mimetic, BAY 85-3934, for the entire course of the experiment. For hypoxia-treatment, cells were treated with intermittent hypoxia. Cells were placed into hypoxic conditions for a total of 48 hours, in 24 hour intervals. (F) T7 endonuclease I assay measuring Cas9-mediated cleavage at the PLK1 locus in HeLa cells 72 hours post addition of LNPs packaged with the indicated constructs and treated in the indicated conditions. Data are represented as mean ± S.D of triplicate treated samples, except (F) which represents mean ± S.D of two independent samples; n.s. not significant; ***p<0.001; ****p<0.0001.

Given our success in driving hypoxia specific gene expression of eGFP, we sought to evaluate the efficacy of delivering HRE promoter driven HSV-tk and Cas9 therapeutic cargo into hypoxic cells using the KC2 LNP formulation. Treatment of tumor cells with our HSV-tk packaged LNPs displayed a significant reduction in cell viability in HRE HSV-tk constructs cultured in GCV with hypoxia mimetic or true hypoxia (Figures 4D and S4E-F). In contrast, little to no change in viability was observed in HeLa cells receiving HRE driven HSV-tk LNPs in the presence of GCV and DMSO. Likewise, we were able to observe a significant reduction in cell viability in a hypoxia regulated manner for cells treated with HRE-Cas9 packaged LNPs (Figure 4E and S4G). Cas9 mediated cleavage was, once again, shown to occur in a hypoxia specific fashion at the PLK1 locus for cells treated with Cas9 expressed from the HRE promoter (Figures 4F and S4H). Overall, these results demonstrate efficient hypoxia regulated expression and activity of therapeutically relevant cargo delivered to cells using plasmid-packaged LNPs.

## Discussion

Hypoxia is present in the majority of solid tumors as a result of alterations in local vascular blood flow that lead to oxygen deprivation. This phenomenon presents a unique opportunity to selectively target the solid tumor microenvironment, without significantly affecting the surrounding normoxic tissue. Multiple studies have shown the benefit of targeting this microenvironment, as hypoxic cells are known to be more resistant to radio- and chemotherapy (3,6). In addition to this, selective killing of HIF1α-expressing tumor cells has been shown to reduce metastatic progression and improve survival (49), suggesting that targeting these cells is a viable therapeutic strategy. In this study, we developed an LNP-based platform to deliver therapeutically relevant cargo to hypoxic tumor cells. We focused our efforts on delivery of two different anti-cancer therapeutic systems: HSV-tk and the spCas9 nuclease. Using 5HRE driven expression, we were able to show LNP-delivered robust and specific expression of these proteins in hypoxic conditions, with minimal off-target activity in normoxic conditions in multiple cancer cell lines. Notably, we were able to reach similar levels of activity when delivering our DNA constructs through LNPs formulated within our lab when compared to commercially available transfection reagents.

Our proof-of-concept work demonstrates an effective delivery of a hypoxia responsive HSV-tk and CRISPR/Cas9 system using nanoparticles; however, further optimization to include additional levels of regulation should be explored. Although we observe a strong induction of gene expression using the 5HRE promoter during hypoxic culture conditions, a small amount of baseline expression persists in control cells. One method to help reduce baseline expression from the 5HRE promoter is to optimize the incorporation of the HIF1α ODD. Installing two layers of regulation with the 5HRE and ODD would greatly limit any residual expression in normoxia which has been demonstrated in CAR-T cells (9). To further tailor the 5HRE system for expression in tumor cells, the inclusion of tumor-specific promoters should be explored. Previous work has demonstrated the use of Survivin, TERT, CEA, PSA, and other gene promoters for cancer specific expression of transgenes (50). The inclusion of a cancer specific promoter would help limit expression of the 5HRE system in normal cells that may have high levels of VEGF, such as hematopoietic stem cells (51). Furthermore, this would allow for the targeting of normoxic tumor cells, while enhancing the targeting of the resistant hypoxic cell population.

Another strategy that we envision could increase expression specificity for our genes of interest in tumor cells is to incorporate microRNA (miRNA) seed sequences into the 3’ untranslated region (UTR) of the gene. miRNAs are small single-stranded RNAs that bind to mRNAs based on sequence complementarity and prevent protein expression by either promoting degradation of the mRNA transcript or blocking translation machinery. This strategy for post-transcriptional gene silencing can be implemented when there is dysregulated expression of miRNAs between the target and non-target cells. Previous studies have capitalized on the dysregulated expression of miR-122 for use in delivering HSV-tk gene for hepatocellular carcinoma (52,53). Beyond this, miR-128 and miR-124 seed sequences have been used to achieve glioblastoma specific expression of transgenes in combination with cancer specific promoters (54,55).

In addition to utilizing methods of controlling gene expression on the transcriptional and post-transcriptional level, advances in LNP delivery and targeting may also be combined with the 5HRE constructs described here. Although most LNPs primarily traffic to the liver, modifications to the surface of the LNP can increase delivery to tumor cells. Addition of an EGFR antibody to the surface of an LNP using Anchored Secondary ScFv Enabling Targeting (ASSET) technology significantly enhanced LNP uptake and delivery of CRISPR/Cas9 mRNA in OV8 ovarian tumors (45). Furthermore, there are stimuli-responsive lipids and peptides that can be incorporated into LNPs to promote penetration within a tumor in order to reach the hypoxic core (45,56).

In conclusion, we provide proof-of-concept work demonstrating the feasibility of targeting HSV-tk and Cas9 therapeutic cargo to hypoxic tumor cells using LNPs. To the best of our knowledge this is the first report demonstrating hypoxia specific expression and activity of the CRISPR/Cas9 nuclease in tumor cells. This is also the first report proposing the delivery of a DNA based hypoxia regulated gene therapy system using LNPs. The technologies presented here hold tremendous potential in targeting some of the most aggressive and treatment resistant cells within a solid tumor and should be explored further for enhanced cancer specificity and delivery.

## Acknowledgements

We thank the City of Hope Core Facilities for their support. In particular, we thank Dr. Zhuo Li and Dr. Ricardo Zerda at the COH Electron Microscopy and Atomic Force Microscopy core, as well as Dr. Brian Armstrong at the COH Light Microscopy/Digital Imaging core for their expertise and technical assistance. We thank the COH Analytical Cytometry core for their assistance.

This work was supported by NIMH R01 113407-01 to KVM and core services at COH supported by the National Cancer Institute of the National Institutes of Health award number P30CA033572.

## Author contributions

G.S. and A.M.D. conceived and designed the project and carried out the experiments. G.S., A.M.D., and K.V.M. drafted the manuscript.

## References

1. Surveillance, Epidemiology, and End Results Program [Internet]. [cited 2021 Jun 22]. Available from: https://seer.cancer.gov/index.html

2. Hematopoietic Cancers. Primer to the Immune Response. Academic Cell; 2014. page 553–85.

3. Vaupel P, Mayer A. Hypoxia in cancer: significance and impact on clinical outcome. Cancer Metastasis Rev. 2007;26:225–39.

4. Bernauer C, Stella Man YK, Chisholm JC, Lepicard EY, Robinson SP, Shipley JM. Hypoxia and its therapeutic possibilities in paediatric cancers. Br J Cancer. Nature Publishing Group; 2020;124:539–51.

5. Höckel M, Vaupel P. Tumor Hypoxia: Definitions and Current Clinical, Biologic, and Molecular Aspects. J Natl Cancer Inst. Oxford Academic; 2001;93:266–76.

6. Dhani N, Fyles A, Hedley D, Milosevic M. The clinical significance of hypoxia in human cancers. Semin Nucl Med. 2015;45:110–21.

7. Huang LE, Gu J, Schau M, Bunn HF. Regulation of hypoxia-inducible factor 1alpha is mediated by an O2-dependent degradation domain via the ubiquitin-proteasome pathway. Proc Natl Acad Sci U S A. 1998;95:7987–92.

8. Iommarini L, Porcelli AM, Gasparre G, Kurelac I. Non-Canonical Mechanisms Regulating Hypoxia-Inducible Factor 1 Alpha in Cancer. Front Oncol [Internet]. Frontiers; 2017 [cited 2021 Jun 22];7. Available from: https://www.frontiersin.org/articles/10.3389/fonc.2017.00286/pdf

9. Kosti P, Opzoomer JW, Larios-Martinez KI, Henley-Smith R, Scudamore CL, Okesola M, et al. Hypoxia-sensing CAR T cells provide safety and efficacy in treating solid tumors. Cell Rep Med. 2021;2:100227.

10. Harada H, Hiraoka M, Kizaka-Kondoh S. Antitumor Effect of TAT-Oxygen-dependent Degradation-Caspase-3 Fusion Protein Specifically Stabilized and Activated in Hypoxic Tumor Cells. Cancer Res. American Association for Cancer Research; 2002;62:2013–8.

11. Harvey TJ, Hennig IM, Shnyder SD, Cooper PA, Ingram N, Hall GD, et al. Adenovirus-mediated hypoxia-targeted gene therapy using HSV thymidine kinase and bacterial nitroreductase prodrug-activating genes in vitro and in vivo. Cancer Gene Ther. 2011;18:773–84.

12. Shibata T, Giaccia AJ, Brown JM. Development of a hypoxia-responsive vector for tumor-specific gene therapy. Gene Ther. 2000;7:493–8.

13. Karjoo Z, Chen X, Hatefi A. Progress and problems with the use of suicide genes for targeted cancer therapy. Adv Drug Deliv Rev. 2016;99:113–28.

14. Westphal M, Ylä-Herttuala S, Martin J, Warnke P, Menei P, Eckland D, et al. Adenovirus-mediated gene therapy with sitimagene ceradenovec followed by intravenous ganciclovir for patients with operable high-grade glioma (ASPECT): a randomised, open-label, phase 3 trial. Lancet Oncol. 2013;14:823–33.

15. Chew WL, Tabebordbar M, Cheng JKW, Mali P, Wu EY, Ng AHM, et al. A multifunctional AAV-CRISPR-Cas9 and its host response. Nat Methods. 2016;13:868–74.

16. Luther DC, Lee YW, Nagaraj H, Scaletti F, Rotello VM. Delivery approaches for CRISPR/Cas9 therapeutics in vivo: advances and challenges. Expert Opin Drug Deliv. 2018;15:905–13.

17. Ronzitti G, Gross D-A, Mingozzi F. Human Immune Responses to Adeno-Associated Virus (AAV) Vectors. Front Immunol. 2020;11:670.

18. Wilson RC, Gilbert LA. The Promise and Challenge of In Vivo Delivery for Genome Therapeutics. ACS Chem Biol. 2018;13:376–82.

19. Mitchell MJ, Billingsley MM, Haley RM, Wechsler ME, Peppas NA, Langer R. Engineering precision nanoparticles for drug delivery. Nat Rev Drug Discov. Nature Publishing Group; 2020;20:101–24.

20. Oberli MA, Reichmuth AM, Robert Dorkin J, Mitchell MJ, Fenton OS, Jaklenec A, et al. Lipid Nanoparticle Assisted mRNA Delivery for Potent Cancer Immunotherapy. Nano Lett. NIH Public Access; 2017;17:1326.

21. Zatsepin TS, Kotelevtsev YV, Koteliansky V. Lipid nanoparticles for targeted siRNA delivery – going from bench to bedside. Int J Nanomedicine. Dove Press; 2016;11:3077.

22. Wei T, Cheng Q, Min Y-L, Olson EN, Siegwart DJ. Systemic nanoparticle delivery of CRISPR-Cas9 ribonucleoproteins for effective tissue specific genome editing. Nat Commun. Nature Publishing Group; 2020;11:1–12.

23. Kulkarni JA, Myhre JL, Chen S, Tam YYC, Danescu A, Richman JM, et al. Design of lipid nanoparticles for in vitro and in vivo delivery of plasmid DNA. Nanomedicine. 2017;13:1377–87.

24. Kulkarni JA, Cullis PR, van der Meel R. Lipid Nanoparticles Enabling Gene Therapies: From Concepts to Clinical Utility. Nucleic Acid Ther. 2018;28:146–57.

25. Sago CD, Lokugamage MP, Paunovska K, Vanover DA, Monaco CM, Shah NN, et al. High-throughput in vivo screen of functional mRNA delivery identifies nanoparticles for endothelial cell gene editing. Proc Natl Acad Sci U S A. National Academy of Sciences; 2018;115:E9944–52.

26. Flamme I, Oehme F, Ellinghaus P, Jeske M, Keldenich J, Thuss U. Mimicking hypoxia to treat anemia: HIF-stabilizer BAY 85-3934 (Molidustat) stimulates erythropoietin production without hypertensive effects. PLoS One. 2014;9:e111838.

27. Leung AKK, Hafez IM, Baoukina S, Belliveau NM, Zhigaltsev IV, Afshinmanesh E, et al. Lipid Nanoparticles Containing siRNA Synthesized by Microfluidic Mixing Exhibit an Electron-Dense Nanostructured Core. J Phys Chem C Nanomater Interfaces. 2012;116:18440–50.

28. Urak RZ, Soemardy C, Ray R, Li S, Shevchenko G, Scott T, et al. Conditionally Replicating Vectors Mobilize Chimeric Antigen Receptors against HIV. Mol Ther Methods Clin Dev. 2020;19:285–94.

29. Jaakkola P, Mole DR, Tian YM, Wilson MI, Gielbert J, Gaskell SJ, et al. Targeting of HIF-alpha to the von Hippel-Lindau ubiquitylation complex by O2-regulated prolyl hydroxylation. Science. 2001;292:468–72.

30. Vordermark D, Shibata T, Brown JM. Green fluorescent protein is a suitable reporter of tumor hypoxia despite an oxygen requirement for chromophore formation. Neoplasia. 2001;3:527–34.

31. Janoly-Dumenil A, Rouvet I, Bleyzac N, Morfin F, Zabot M-T, Tod M. A pharmacodynamic model of ganciclovir antiviral effect and toxicity for lymphoblastoid cells suggests a new dosing regimen to treat cytomegalovirus infection. Antimicrob Agents Chemother. 2012;56:3732–8.

32. Kim YG, Bi W, Feliciano ES, Drake RR, Stambrook PJ. Ganciclovir-mediated cell killing and bystander effect is enhanced in cells with two copies of the herpes simplex virus thymidine kinase gene. Cancer Gene Ther. 2000;7:240–6.

33. Srivastava D, Joshi G, Somasundaram K, Mulherkar R. Mode of cell death associated with adenovirus-mediated suicide gene therapy in HNSCC tumor model. Anticancer Res. 2011;31:3851–7.

34. Howard BD, Boenicke L, Schniewind B, Henne-Bruns D, Kalthoff H. Transduction of human pancreatic tumor cells with vesicular stomatitis virus G-pseudotyped retroviral vectors containing a herpes simplex virus thymidine kinase mutant gene enhances bystander effects and sensitivity to ganciclovir. Cancer Gene Ther. 2000;7:927–38.

35. Ardiani A, Sanchez-Bonilla M, Black ME. Fusion enzymes containing HSV-1 thymidine kinase mutants and guanylate kinase enhance prodrug sensitivity in vitro and in vivo. Cancer Gene Ther. 2010;17:86–96.

36. Kokoris MS, Sabo P, Adman ET, Black ME. Enhancement of tumor ablation by a selected HSV-1 thymidine kinase mutant. Gene Ther [Internet]. Gene Ther; 1999 [cited 2021 Aug 10];6. Available from: https://pubmed.ncbi.nlm.nih.gov/10467366/

37. Martinez-Lage M, Torres-Ruiz R, Puig-Serra P, Moreno-Gaona P, Martin MC, Moya FJ, et al. In vivo CRISPR/Cas9 targeting of fusion oncogenes for selective elimination of cancer cells. Nat Commun. Nature Publishing Group; 2020;11:1–14.

38. Gao Q, Ouyang W, Kang B, Han X, Xiong Y, Ding R, et al. Selective targeting of the oncogenic G12S mutant allele by CRISPR/Cas9 induces efficient tumor regression. Theranostics. 2020;10:5137–53.

39. Yoshiba T, Saga Y, Urabe M, Uchibori R, Matsubara S, Fujiwara H, et al. CRISPR/Cas9-mediated cervical cancer treatment targeting human papillomavirus E6. Oncol Lett. 2019;17:2197–206.

40. Lee S, Kim Y-Y, Ahn HJ. Systemic delivery of CRISPR/Cas9 to hepatic tumors for cancer treatment using altered tropism of lentiviral vector [Internet]. Biomaterials. 2021. page 120793. Available from: http://dx.doi.org/10.1016/j.biomaterials.2021.120793

41. Zhang X-H, Tee LY, Wang X-G, Huang Q-S, Yang S-H. Off-target Effects in CRISPR/Cas9-mediated Genome Engineering. Mol Ther Nucleic Acids. 2015;4:e264.

42. Höijer I, Johansson J, Gudmundsson S, Chin C-S, Bunikis I, Häggqvist S, et al. Amplification-free long-read sequencing reveals unforeseen CRISPR-Cas9 off-target activity. Genome Biol. BioMed Central; 2020;21:1–19.

43. Naeem M, Majeed S, Hoque MZ, Ahmad I. Latest Developed Strategies to Minimize the Off-Target Effects in CRISPR-Cas-Mediated Genome Editing. Cells [Internet]. 2020;9. Available from: http://dx.doi.org/10.3390/cells9071608

44. Weiß L, Efferth T. Polo-like kinase 1 as target for cancer therapy. Exp Hematol Oncol. 2012;1:38.

45. Rosenblum D, Gutkin A, Kedmi R, Ramishetti S, Veiga N, Jacobi AM, et al. CRISPR-Cas9 genome editing using targeted lipid nanoparticles for cancer therapy. Sci Adv [Internet]. 2020;6. Available from: http://dx.doi.org/10.1126/sciadv.abc9450

46. Milane L, Amiji M. Clinical approval of nanotechnology-based SARS-CoV-2 mRNA vaccines: impact on translational nanomedicine. Drug Deliv Transl Res. 2021;11:1309–15.

47. Witzigmann D, Kulkarni JA, Leung J, Chen S, Cullis PR, van der Meel R. Lipid nanoparticle technology for therapeutic gene regulation in the liver. Adv Drug Deliv Rev. 2020;159:344–63.

48. Kulkarni JA, Witzigmann D, Leung J, Tam YYC, Cullis PR. On the role of helper lipids in lipid nanoparticle formulations of siRNA. Nanoscale. The Royal Society of Chemistry; 2019;11:21733–9.

49. Kizaka-Kondoh S, Itasaka S, Zeng L, Tanaka S, Zhao T, Takahashi Y, et al. Selective killing of hypoxia-inducible factor-1-active cells improves survival in a mouse model of invasive and metastatic pancreatic cancer. Clin Cancer Res. 2009;15:3433–41.

50. Robson T, Hirst DG. Transcriptional Targeting in Cancer Gene Therapy. J Biomed Biotechnol. 2003;2003:110–37.

51. Gerber H-P, Malik AK, Solar GP, Sherman D, Liang XH, Meng G, et al. VEGF regulates haematopoietic stem cell survival by an internal autocrine loop mechanism. Nature. 2002;417:954–8.

52. Dhungel B, Ramlogan-Steel CA, Layton CJ, Steel JC. miRNA122a regulation of gene therapy vectors targeting hepatocellular cancer stem cells. Oncotarget. 2018;9:23577–88.

53. Wang G, Dong X, Tian W, Lu Y, Hu J, Liu Y, et al. Evaluation of miR-122-regulated suicide gene therapy for hepatocellular carcinoma in an orthotopic mouse model. Chin J Cancer Res. 2013;25:646–55.

54. Skalsky RL, Cullen BR. Reduced expression of brain-enriched microRNAs in glioblastomas permits targeted regulation of a cell death gene. PLoS One. 2011;6:e24248.

55. Wu C, Lin J, Hong M, Choudhury Y, Balani P, Leung D, et al. Combinatorial control of suicide gene expression by tissue-specific promoter and microRNA regulation for cancer therapy. Mol Ther. 2009;17:2058–66.

56. Kulkarni P, Haldar MK, Katti P, Dawes C, You S, Choi Y, et al. Hypoxia Responsive, Tumor Penetrating Lipid Nanoparticles for Delivery of Chemotherapeutics to Pancreatic Cancer Cell Spheroids. Bioconjug Chem. 2016;27:1830–8.

