## Supplemental Figures for "Hypoxia-directed tumor targeting of CRISPR/Cas9 and HSV-TK suicide gene therapy using lipid nanoparticles"

### SUPPLEMENTAL DATA FIGURES

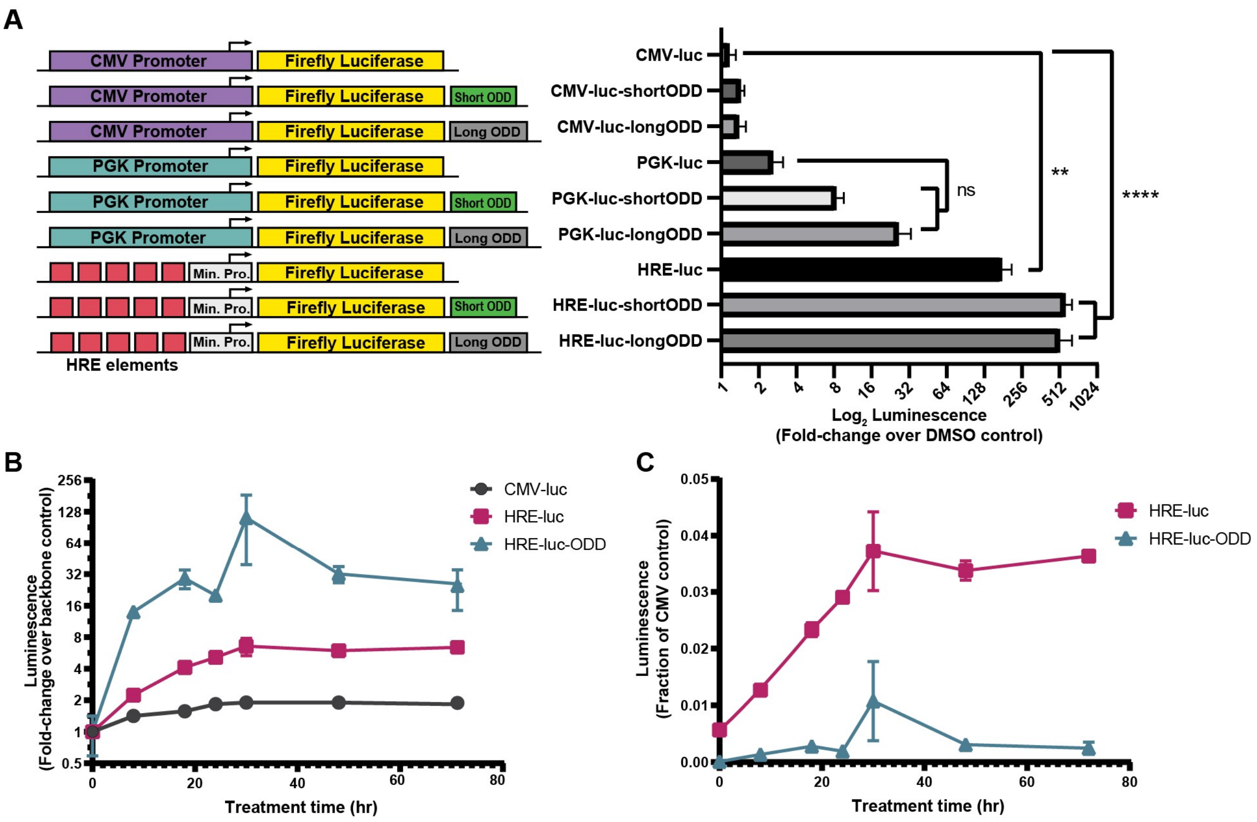


**Supplementary Figure 1**

(A) Relative increase in luciferase activity for the indicated constructs in HeLa cells treated with 10uM BAY 85-3934 for 12 hours, compared to the DMSO-treated (0.1% v/v DMSO) control. (B) Time-course induction of normalized luciferase activity relative to the respective backbone control in NCI-H1299 cells expressing the indicated constructs grown in the presence of 10µM BAY 85-3934, or vehicle control (0.1% v/v DMSO). (C) Same as in (B), except the luciferase signal is normalized to the CMV backbone. Data are represented as mean ± S.D of triplicate treated samples; n.s. not significant; **p<0.01; ****p<0.0001.

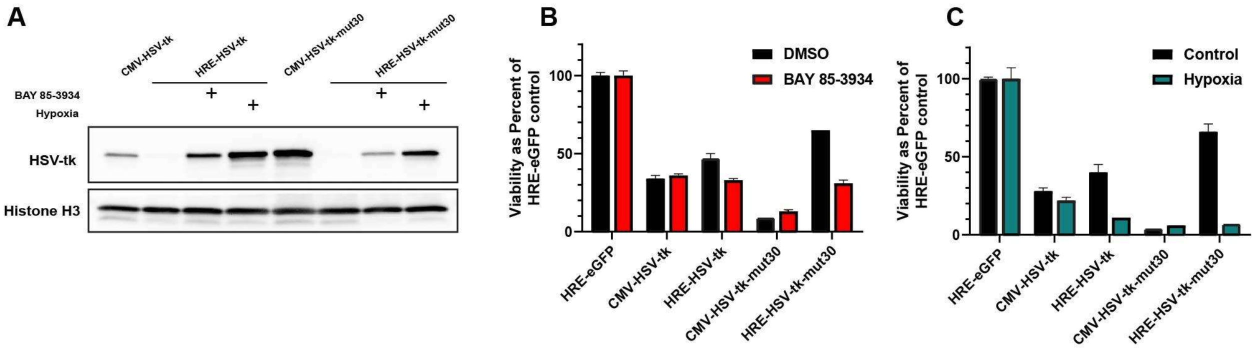


**Supplementary Figure 2**

(A) Western blot analysis of U251 cells transfected with either the CMV or HRE-driven HSV-tk WT or mutant 30 constructs. Transfected cells were cultured in the presence of 10 μM BAY 85-3934 or hypoxia for 24 hours prior to harvesting for protein analysis. (B) An Alamar assay for viability was carried out at 120 hours for U251 cells transfected with HSV-tk constructs and cultured in 2μM GCV and 10μM BAY 85-3934. (C) An Alamar assay for viability was carried out at 120 hours for U251 cells transfected with indicated HSV-tk constructs and treated with 2μM GCV and hypoxia. Cells undergoing hypoxia treatment were exposed to hypoxia for 24 hours during two intermittent periods. Data are represented as mean ± S.D of triplicate treated samples.

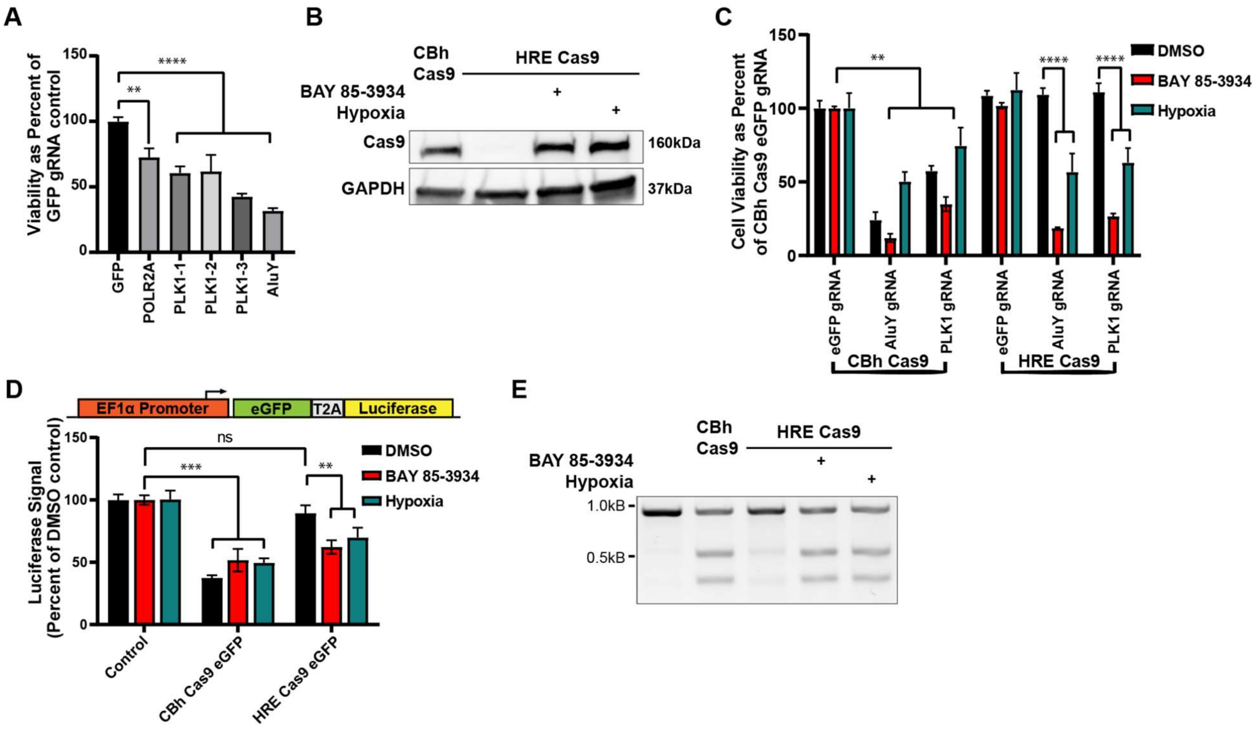


**Supplementary Figure 3**

(A) Alamar assay depicting viability of HEK293 cells 72 hours post-transfection with a CBh-driven Cas9 and the indicated gRNAs. (B) Western blot showing expression of Cas9 in U251 cells transfected with either the CBh- or HRE-driven Cas9. Cells transfected with the HRE-driven Cas9 were cultured in the presence of 10µM BAY 85-3934 or hypoxia for 24 hours before harvesting lysates. (C) Alamar assay comparing viability of U251 cells 120 hours post-transfection with the indicated constructs. Cells were cultured in the presence of the hypoxia mimetic, BAY 85-3934, for the entire course of the experiment. For hypoxia-treatment, cells were treated with intermittent hypoxia. Cells were placed into hypoxic conditions for a total of 48 hours, in 24 hour intervals. (D) HeLa cells were stably transduced with an eGFP-firefly luciferase construct driven from an EF1α promoter, as indicated in the schematic at the top. After generation of the stable line, cells were transfected with the indicated constructs and relative luciferase activity was measured after 48 hours. (E) Representative 2% agarose gel image of T7 endonuclease mediated cleavage at the PLK1 locus in HeLa cells 48 hours post-transfection with the indicated constructs. Data are represented as mean ± S.D of triplicate treated samples; n.s. not significant; **$p<0.01$; ***$p<0.001$; ****$p<0.0001$.

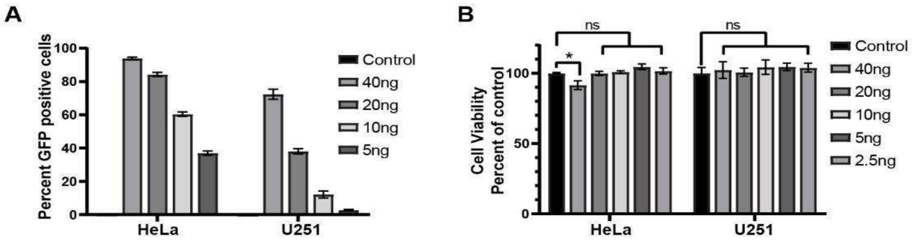


C

| KC2 DNA-LNP | Size (nm) | PDI | Zeta potential (mV) | EE% |
|---|---|---|---|---|
| HRE-eGFP | 133.7±0.7 | 0.068±0.008 | 15.37±1.34 | 66.1 |
| Cas9-T2A eGFP | 149.6±1.6 | 0.117±0.011 | 21±1.55 | 76.4 |
| HRE HSV-tk WT | 122.7±0.3 | 0.084±0.007 | 22.5±1.49 | 73.3 |
| CMV HSV-tk WT | 148.5±1.7 | 0.101±0.014 | 21.68±0.67 | 44.2 |
| HRE Mutant 30 | 122.6±1.1 | 0.08±0.017 | 27.45±1.76 | 75.8 |
| CMV Mutant 30 | 119±0.9 | 0.061±0.006 | 16.64±0.83 | 75.5 |
| HRE BbsI Cas9 | 133.9±1.1 | 0.078±0.026 | 22.78±1.64 | 66.3 |
| Cbh BbsI Cas9 | 130.7±0.9 | 0.052±0.011 | 22.68±1.78 | 60.6 |
| HRE AluY Cas9 | 140.6±1.4 | 0.075±0.012 | 19.73±1.6 | 75.1 |
| Cbh AluY Cas9 | 141.2±1.1 | 0.071±0.015 | 15.34±0.94 | 68.8 |
| HRE PLK Cas9 | 145.2±1.3 | 0.104±0.01 | 35.48±1.05 | 75.3 |
| Cbh PLK Cas9 | 146±1.3 | 0.09±0.015 | 15.95±1.13 | 69.8 |

D

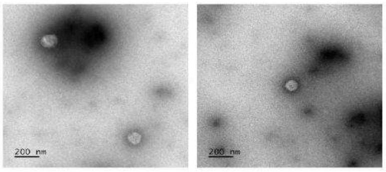

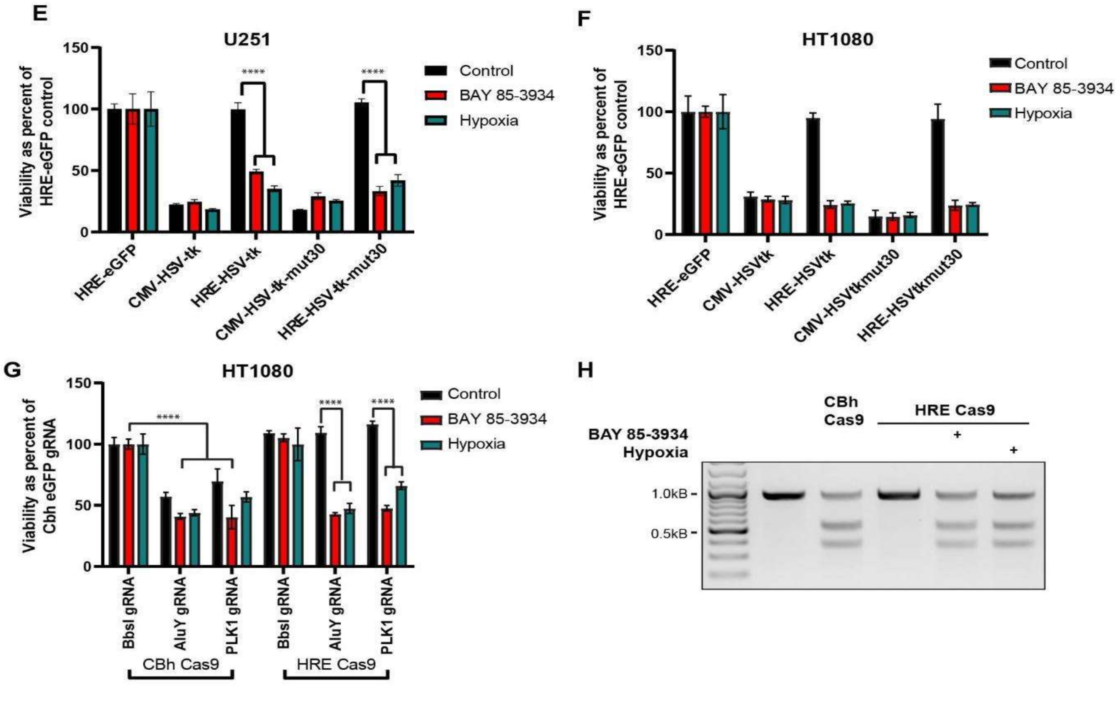


**Supplementary Figure 4**

(A) Flow cytometry analysis of eGFP expression in HeLa and U251 cells 72 hours after addition of the indicated doses of LNPs packaged with the Cas9-T2A-eGFP plasmid. (B) Alamar assay evaluating HeLa and U251 cell viability 48 hours after treatment with increasing concentrations of Cas9-eGFP LNP ranging from 5ng to 40ng of encapsulated DNA. (C) Full physical characterization of size, polydispersity (PDI), zeta potential, and encapsulation efficiency (EE%) for KC2 DNA-LNPs used in this study. Data represented as mean ± S.E.M. based on the number of repeated measurements taken (size and polydispersity=5 runs, zeta potential=10 runs). (D) Representative transmission electron microscopy images of KC2 DNA-LNPs. Scale bar= 200 nm. (E) Alamar assay evaluating potency of LNPs delivering the indicated HSV-tk constructs to U251 cells in the presence of 1µM GCV and DMSO, BAY 85-3934, or hypoxia treatment. Viability was measured at 120 hours post LNP treatment. (F) same as in (E) except for HT1080 cells. (G) Alamar assay comparing viability of HT1080 cells 120 hours post addition of LNPs packaged with the indicated constructs. Cells were cultured in the presence of the hypoxia mimetic, BAY 85-3934, for the entire course of the experiment. For hypoxia-treatment, cells were treated with hypoxia for 24 hours. (H) Representative 2% agarose gel image of T7 endonuclease mediated cleavage at the PLK1 locus in HeLa cells 72 hours post addition of LNPs packaged with the indicated constructs and treated in the indicated conditions. Data are represented as mean ± S.D of triplicate treated samples; n.s. not significant; *$p<0.05$; ****$p<0.0001$.
