## Supplemental Table for "Hypoxia-directed tumor targeting of CRISPR/Cas9 and HSV-TK suicide gene therapy using lipid nanoparticles"

### Supplemental Table 1: List of Plasmids used in this study

| Name | Source | Information |
|---|---|---|
| 5HRE/GFP | Addgene 46926 | Used to PCR out the 5xHRE promoter using oligos HRE1-2 and cloned using NEBuilder® HiFi DNA Assembly into JDS246 (NotI/MluI digested) and pcDNA3-luciferase (HindIII/MluI digested) |
| pRRlsinPGK_BFP2_WPRE | Addgene 91787 | Used to PCR out the hPGK promoter using oligos PGK1-2 and cloned using NEBuilder® HiFi DNA Assembly into JDS246 (NotI/MluI digested) and pcDNA3-luciferase (HindIII/MluI digested) |
| pcDNA3-ffluc | This lab | Expresses firefly luciferase from a CMV promoter. |
| pcDNA3-ffluc-shortODD | This lab | Expresses firefly luciferase short ODD fusion protein driven from a CMV promoter. |
| pcDNA3-ffluc-longODD | This lab | Expresses firefly luciferase long ODD fusion protein driven from a CMV promoter. |
| PGK-ffluc | This lab | Expresses firefly luciferase from a PGK promoter. |
| PGK-ffluc-shortODD | This lab | Expresses firefly luciferase short ODD fusion protein driven from a PGK promoter. |
| PGK-ffluc-longODD | This lab | Expresses firefly luciferase long ODD fusion protein driven from a PGK promoter. |
| 5HRE-ffluc | This lab | Expresses firefly luciferase from a 5HRE promoter. |
| 5HRE-ffluc-shortODD | This lab | Expresses firefly luciferase short ODD fusion protein driven from a 5HRE promoter. |
| 5HRE-ffluc-longODD | This lab | Expresses firefly luciferase long ODD fusion protein driven from a 5HRE promoter. |
| pAL119-TK | Addgene 21911 | CMV driven HSV-TK |
| CMV-HSV-tk-mut30 | This lab | Expresses HSV-tk mutant 30 (L159I, I160L, F161A, A168Y, L169F) from a CMV promoter |
| 5HRE-HSV-tk | This lab | 5HRE driven HSV-TK |
| 5HRE-HSV-tk-mut30 | This lab | Expresses HSV-tk mutant 30 (L159I, I160L, F161A, A168Y, L169F) from a 5HRE promoter |
| 3XFLAG-Cas9 | This lab | Expresses Cas9 nuclease with N-terminal 3X FLAG from CBh promoter (CMV early enhancer; ch-beta actin promoter; chimeric ch-beta actin MVM intron). Does not contain any gRNA or fluorescent tag. |
| 5HRE-Cas9 | This lab | 5HRE driven expression of spCas9. Does not contain any gRNA or fluorescent tag. |
| eGFP gRNA | This lab | Expresses "GACGTAGCCTTCGGGCATGG" gRNA for spCas9 targeting eGFP from a H1 promoter |
| AluY gRNA | This lab | Expresses "ACCTGTAGTCCCAGCTACTC " gRNA for spCas9 targeting AluY repeats from a H1 promoter |
| PLK1 gRNA-1 | This lab | Expresses "ACCTCGGGAGCTATGTAATT " gRNA for spCas9 targeting PLK1 gene from a H1 promoter |
| PLK1 gRNA-2 | This lab | Expresses "CGACTTCGTGTTCGTGGTGT " gRNA for spCas9 targeting PLK1 gene from a H1 promoter |
| PLK1 gRNA-3 | This lab | Expresses "TGCTGCTCAAGCCGCACCAG " gRNA for spCas9 targeting PLK1 gene from a H1 promoter. Best candidate that was used in all Cas9 experiments in Fig. 3-4 |
| CBh-Cas9 BbsI gRNA | This lab | Expresses Cas9 from a CBh promoter, in addition to "GACGTAGCCTTCGGGCATGG" gRNA for spCas9 targeting eGFP from a U6 promoter |
| CBh-Cas9 AluY gRNA | This lab | Expresses Cas9 from a CBh promoter, in addition to "ACCTGTAGTCCCAGCTACTC " gRNA for spCas9 targeting AluY repeats from a U6 promoter |
| CBh-Cas9 PLK1 gRNA | This lab | Expresses Cas9 from a CBh promoter, in addition to "TGCTGCTCAAGCCGCACCAG " gRNA for spCas9 targeting PLK1 gene from a U6 promoter |
| HRE-Cas9 BbsI gRNA | This lab | Expresses Cas9 from an HRE promoter, in addition to "GACGTAGCCTTCGGGCATGG" gRNA for spCas9 targeting eGFP from a U6 promoter |
| HRE-Cas9 AluY gRNA | This lab | Expresses Cas9 from an HRE promoter, in addition to "ACCTGTAGTCCCAGCTACTC " gRNA for spCas9 targeting AluY repeats from a U6 promoter |
| HRE-Cas9 PLK1 gRNA | This lab | Expresses Cas9 from an HRE promoter, in addition to "TGCTGCTCAAGCCGCACCAG " gRNA for spCas9 targeting PLK1 gene from a U6 promoter |

| | | |
|---|---|---|
| pLenti-EF1a-GFP-ffluc-BSR | This lab | Third generation lentiviral packaging construct expressing GFP, ffluc and blasticidin resistance gene from an EF1a promoter |
| pMDLg/pRRE | Addgene 12251 | 3rd generation lentiviral packaging plasmid; Contains Gag and Pol |
| pRSV-Rev | Addgene 12253 | 3rd generation lentiviral packaging plasmid; Contains Rev |
| pMD2.G | Addgene 12259 | VSV-G envelope expressing plasmid |
